# Modeling and Visualization of Rice Root Based on Morphological Parameters

**DOI:** 10.1101/553925

**Authors:** Le Yang, Peng Shao

## Abstract

To clarify the morphological distributional characteristic of rice roots, the “root box” experiments are conducted to extract various morphological parameters of roots. On the basis of experiments, in this paper, the rice root model based on morphological parameters is constructed with B-spline curves by analyzing the topological structure of rice roots, quantifying their biological characteristics, summarizing the morphological structure and growth characteristics and improving the Cubic growth function to describe the growth change of rice roots. Meanwhile, the output accuracy of the model is tested. Finally, the dynamic simulation of rice root growth characteristics in three-dimensional space is implemented by using Visual C++ and OpenGL standard graphics library. The compared results demonstrate that the model could faithfully simulate the dynamic growing process of rice roots, and help to enrich the methods of digitization and visualization for roots of other crops.

## 1 Introduction

The functions of the plant roots are mainly used to absorb moisture and nutrient ions in the surrounding soil (Kreszies 2018). As the “Engine” of rice growth, roots absorb water and nutrient needed by plant growth and their morphological and physiological indexes are closely related to the prophase growing development of the overground part of roots, the crop output and grain quality in the later stage. Therefore, it is the basis and guarantees to maintain the crop yield for the better root morphological and physiological characteristics (Somaweera 2017). However, around 75 % of rice is cultivated in paddy soils under flooding condition during the growing season (Atulba 2015). All important part of plants is typically hidden and difficult to measure and study (Dunbabin 2013).Moreover, the rice roots have no more prominent growth characteristics similar to main stems and leaves such as nodes. Therefore, it is more difficult to quantify the structure of rice roots such as partitioning of basic units. The root research is an important part of the rice science. The first descriptive threedimensional root system models were presented by Pages et al. (Pages 1989) and Diggle (Diggle 1988). New root growth models with a strong focus on visualization were developed (Spek 1997 and Lynch 1997). Pages et al. (Pages 2004) proposed the RootTyp model which is a general model of roots. The RootTyp model describes systematically the whole process of growing development of roots including the growth position of roots, axial growth and radial growth direction and root branching and so on. It implements the simulation of different roots. Dusserre et al. (Dusserre 2009) proposed a simple generic model for upland rice root length density estimation from root intersections on soil profile. Leitner et al. (Leitner 2010) introduced a parametric L-System model for root system growth. It is showed that the model can reproduce well-known illustrations of three topologically different root systems. Leitner et al. (Leitner 2010) adopted L-system to describe the interaction between roots and soil, and take into consideration the effects of various tropic movements on the root growth. Ge et al. (Ge 2011) established a three-dimensional visual simulation model by using architectural parameters of upland rice root system. Clark et al. (Clark 2011) introduced a novel 3D imaging and software platform for the high-throughput phenotyping of 3-dimensional root traits during seedling development. Zheng et al. (Zheng 2012) constructed the morphological structure model of rice root using the combination of dual-scale automaton and L-system. Yang et al. (Yang 2011, 2017) built the mathematical models of morphological development of rice roots using crop simulation technologies by analyzing the change pattern of the rice root growth along with the process of development on the basis of analyzing the growth development rule of rice roots. Han et al. (Han 2018) constructed the 3D images of the rice roots, and the phenotypic traits were quantified from the 3D images.

The solving problems, which these research achievements mentioned above, are the morphology and physiological functions based on rice roots, the relationship of roots and over ground parts of roots, and the effects of cultivation and management measures over roots. But they did not consider the combination of the morphological structure and model parameters. In addition, the image simulation results were not ideal and the system operation platform for the human-machine interface was not developed. To observe and analyze accurately the growth and distribution characteristics of rice roots in underground space, this paper adopts the “root box” to make experiments and utilizes the root special scanner and morphological special analysis software to achieve the correlation of root morphological parameters such as the total root length, total root area and root volume of rice roots. In this paper, the optimal regression model, which is between morphological indices of rice roots, is fitted by analyzing the mutation characteristics of root morphological indices. The visualization simulation of rice roots is implemented combined the cubic growth function and B-spline curve with L-system to simulate the curve rice roots with root nodes as the center under the environment of the Visual C++ combined with the OpenGL standard graphics library.

## 2 Materials and methods

### 2.1 The experiment design

The “root box” is adopted to do experiments in the experimental station of Jiangxi Agriculture University during 2016 to 2018. The Ganxin 203 is used as an experimental object to make the box planting experiment with its length, width and height which are all 40cm. The boxes are separated into some small cells with 20 cells in each floor by the stainless steel wire mesh and a volume of a cell is 0.1 m^3^. The “root box” is entangled by the nylon fiber which water and fertilizer can pass but roots cannot pass freely to ensure the integrity of rice root morphology, as shown in Figure 1.

**Fig.1.**
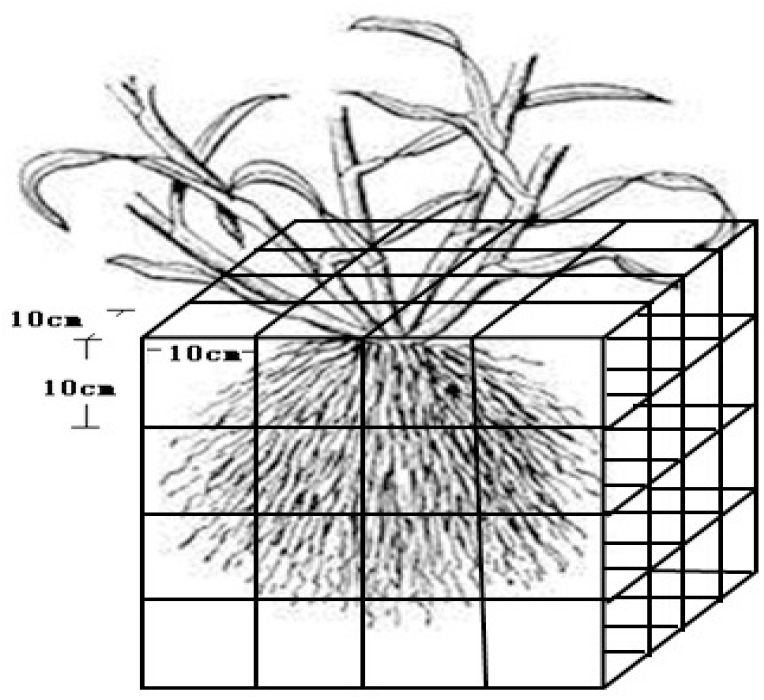
Schematic sketch of the “root box”

The planting method is as follows. Firstly, several mud holes are dug in soil and the “root box” is placed into the mud holes before transplanting rice seedlings. Next, the soil is taken layer by layer with 0.1m as the unit which is buried in the “root box” and it should pay attention to maintaining the original soil composition and the weight of the container. In green houses of the laboratory, rice seedlings are cultivated by the way of floating seedlings (Punsalan 2013) and after 60 days of growth and development the rice seedlings are transplanted into the “root box” of which row-plant space is 20cm ×15cm and planting aperture is 4.5cm. After these operations, the rice nutrient solution is added to the “root box” to meet the normal growth needs of them.

### 2.2 Measure projects and simulation methods

Before taking the rice roots, the “root box” is immersed fully in the 100g/L NaCl solution. It is washed with tap water from top to bottom to obtain the whole rice roots three hours later. Then the root soil of rice is cleaned carefully layer by layer. While the whole rice roots are exposed on the surface part, it is time to observe and analyze the morphological characteristics of rice roots and occurrence of branching roots. Meanwhile, the position and growth direction of each single root are recorded one by one.The morphological parameters such as the root length, diameter and thickness of rice roots are measured. Then the average root length, the total root area and the total root volume are calculated and analyzed. According to the experimental data of the rice morphological features, the visualization simulation of rice roots is implemented combined B-spline curve with L-system by using L-system to generate self-similar grammatical rules for fractal reconstruction of rice roots.

## 3 Construction of rice root models

### 3.1 The morphological characteristics of rice roots

The rice roots belong to the fibrous roots, which can be divided to two categories of the seminal roots and adventitious roots (Ackermann 2013). But only one seminal root in the roots plays functions of absorbing water and nutrition during the rice seedling growth and development. The main part of the roots is adventitious roots which are produced from the residual part of rice stems. The branch roots produced from the adventitious roots of rice roots are called as once branch roots. The branch roots produced from the once branch roots are called as the secondary branch roots. The number of the branch roots can be generated from five to six times under the condition of high yield of rice. Their morphological structure is with the characteristic of self-similarity. During the growth and development of rice continuously, the number of the adventitious roots rooting is also changing correspondingly. In the rice seedlings stage, the number of the adventitious roots rooting is not large but along with the continuous promotion of rice growth and development the number of the adventitious roots is increasing day by day. When the rice is heading nearly, their roots are the most prosperous and the branch roots of all levels are growing in a crisscross pattern. Therefore, the dense and powerful root structure has been formed. The formal diagrammatic sketch of rice roots is shown in Figure 2.

**Fig.2.**
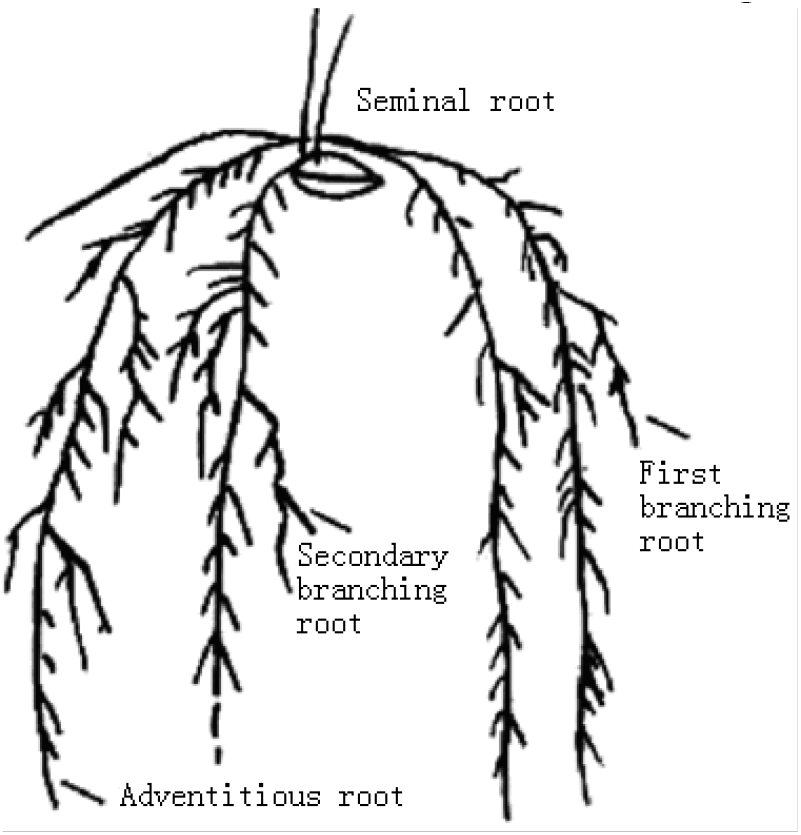
Root architecture of rice

### 3.2 Selection and improvement of the growth functions

What the growth function of rice roots describes is that growth factors (such as the growth length of seminal roots, the number of branch roots of adventitious roots producing) produce the continuous process along with the time change in the condition of nature growth. The growth function unifies its parameters in the form of expressions. The growth function can unifies its parameters with expressions which are obtained by the functional expression of experimental data achieved usually from the growth and development of rice roots directly which are analyzed by mathematical statistics. There is obvious self-similarity between main roots and branching roots according to the growth rhythm of rice roots. Thus the Cubic growth function is chosen to describe the growth change of roots, which is expressed as formulas (1).

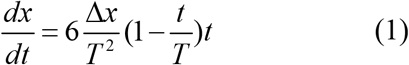

Where the parameter *T* is a growth cycle and *t* represents the time between 0 and *T*. The initiate value *x*_0_ of the root growth increases to the growth terminal value *x_max_* continuously which is used to describe the cubic equation of the change process and meets *x*’ (0)=*x*’ (*T*)=0 and Δ *x*=*x_max_ - x*_0_.

However, based on the real growth situation of rice roots, we can analyze that the growth rate of roots are different even at the same time and the same plant, which is obviously contrary to the natural law when the same growth function is used. For solving this disadvantage, the parameter *rnd*(1) is added into the Cubic growth function as the regulation and control of growth rate The function definition domain is defined at the growth cycle of geometric parameters, which is simple and intuitive. The improved growth function not only maintains the growth characteristics of rice roots, but also regulates and controls the growth rate by simple parameter *rnd*(1) to solve the problem of speed difference between the above roots. The improved function form is expressed as formulas (2).

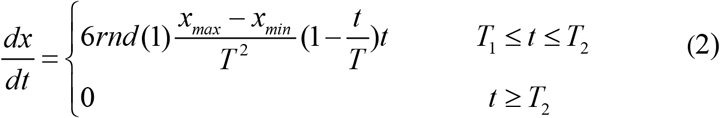

Where the parameter *x_max_* represents the maximum of geometric parameters and the parameter *x_min_* represents the minimum of geometric parameters for root growth. The growth cycle *T* equals *T*_2_ – *T*_1_. The *rnd*(1) is the parameter of speed control and its range is (0, 1].

Compared with the original growth function, the life cycle of the improved function is its definition domain, which makes the expression more intuitive and readable. What is more important is that the difference in growth rate between roots is more consistent with natural growth situation by setting the value of *rnd*(1).

### 3.3 The design of L-system based on B-spline curves

L-systems use grammars which contain a set of rules called production rules, They allow the user to model complex objects by successively replacing parts of a simple object (Bohl 2015).According to the growth characteristics of rice roots, the topological structure of rice roots with self-similarity can be described with the generation rules of L-system. This paper implements the virtual simulation image of roots by adopting the method of the fractal reconstructing morphological structure of rice roots. The B-spline curves are introduced to improve the expression of L-system due to its advantages such as the smoothness, convexity and controllability to express the curve shape of rice roots. In order to gain more local control and admit more complexity of the occluded curve (Zhao 2013), the B-spline curves express the curve roots by the start nodes, the end nodes, the control nodes and the de Boor-box recursive formulas mainly. In the L-system of rice roots, firstly the coordinate of the start nodes and the end nodes representing the character F between the root segments are calculated. Then the control nodes are determined. The control nodes can be added between the start nodes and the end nodes, or be set according to the convex hull direction and the growth space direction of rice roots. Finally, the curve characteristics of rice roots are simulated by de Boor-Cox recursive formulas. The L-system designed based on B-spline curves is shown as formulas (3).

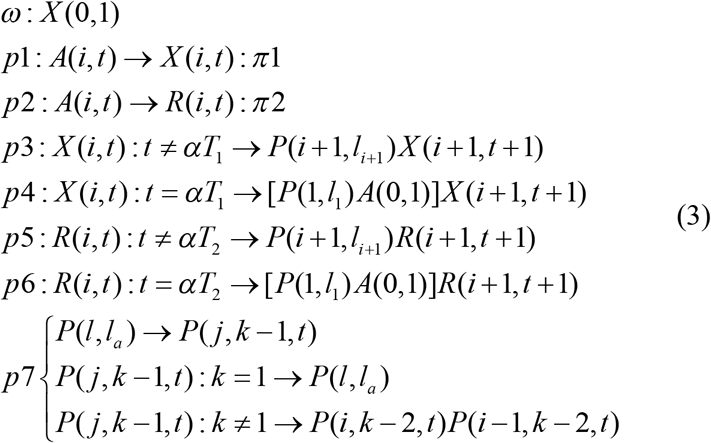

In the L-system of rice roots, the parameter *ω* represents the initial axiom. is the initial strings to express the initial axiom. The growth nodes of roots and *X*(0,1) the growth parameter of nodes are represented with the parameter *p* and *i* respectively.

The parameter α represents the growth direction of roots.The growth time intervals of branch roots of roots use is represented the parameter *t*. The probabilistic values that the system calls production formulas *p*1 and *p*2 are *π*1 and *π*2 respectively. The *A*(*i,t*) represents the growth of branch roots of roots with different probabilities *π*1 and *π*2. The parameter *l* represents the length of branch roots of roots described the growth function. The start node *P*(0) and the end node *P*(*n*) are used to express a curve branch root and *n*-1 control node vector are added between the start nodes and the end nodes according to the direction of the convex hull of roots. Firstly, the initiate position *P*(1,*l*_1_) is generated by the production formula *p*_3_.=The roots produce the first branching roots and secondary branching roots after a growth time interval *t*. *X*(*i, t*) and *R*(*i, t*) represent the different growth points respectively. The growth process of them are represented by the different probabilities *π*1 and *π*2 of *A*(*i, t*).The production formula *p*1 represents the rules of roots and the *P*(*t*) represents the root curve. The parameter *k* represents the order of B-spline and the range of the parameter *i* is 0 ≤ *i* < *n*. The production formula *p*2 and *p*3 are calculated according to the de Boor-Cox recursive formula. If *i* = *j*, *p*2 is called or if *i* = *j* − *k* + *r* + 1’ *p*3 is called, where the parameter *r* is the variable of B-spline curve times. The p7 should be combined with the recursive calculation of b-spline curve, namely formula (4). In order to express the curvature degree of rice roots vividly, it can be expressed by formulas (5) combined with B-spline curves.

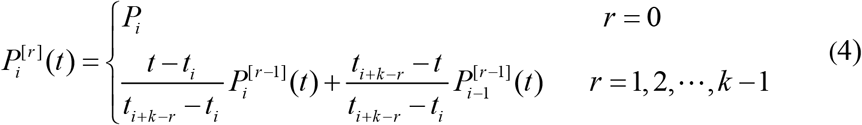

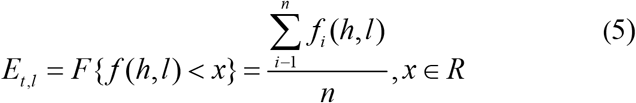

Where the parameters *h* and *l* are the quantification parameters and the parameter *x* is the degree of maximal curve. The *E* and *F* represent the curve line or linear line between two growth points.

## 4 Results and discussion

### 4.1 Model checking

#### (1) Methods of Model checking

To assess the level of agreement between the model-predicted and field-observed data (Dash 2015), correlation coefficient and variation coefficient are combined to test the accuracy of the rice root model.

##### 1) Correlation coefficient

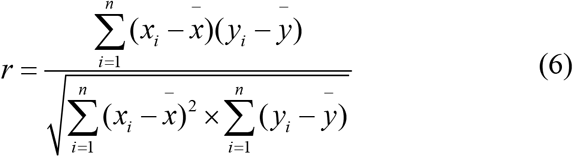

In the formulas (6), the parameter *r* is the correlation coefficient of the model. The parameter *i* and *x_i_* represents the serial number of samples and measured values respectively. The parameter 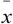 is the average values of measured values. The parameters *y_i_* and *ȳ* are the simulation values and the average values of simulation values respectively. The parameter *n* represents the number of samples.

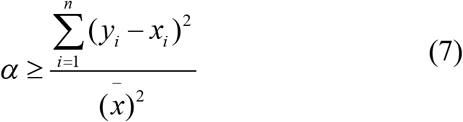

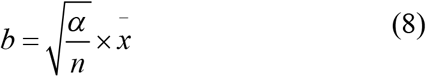

In the formulas (6) and formulas (7), the parameter *α* represents the confidence level of the model and the band domain between 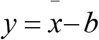, and 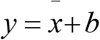, represents the confidence band of *α*.

##### 2) Variation coefficient

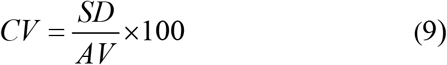

In the formulas (9), the parameter *CV* is the variation coefficient of the model and *AV* is average values and *SD* represents the standard deviation.

#### (2) Comparison of Measure values and simulation values of roots

In the experimental station of Jiangxi Agriculture University, the Ganxin 203 is used as an experimental object. From June 10, 2017, before tillering, the total root length, diameter, total root surface area and total root volume are measured once every two days. When the number of roots is 60 to 100, they are measured twice a week and then once a week. Three plants are taken at each leaf age before the 10^th^ leaf age and then two plants are taken at each leaf age. The root morphology is an important characteristic of the differences of rice varieties. The greater the coefficient of variation (Chittoori 2013), the greater the differences of rice varieties. In this paper, the variation coefficient range of morphological index of rice roots is 0.64 to 0.79, and the latter is 1.23 times of the former. The variation coefficient appears a trend of total root length > total root area > total root volume, which is described in Table 1.

**Table 1.** The morphological parameters of rice seedling roots

| Morphological parameters | Changing range | Average values | CV |
| --- | --- | --- | --- |
| Total root length (cm) | 113.35~2198.22 | 667.61 | 0.79 |
| Total root area (cm<sup>2</sup>) | 14.09~145.14 | 57.11 | 0.67 |
| Total root volume (cm<sup>3</sup>) | 0.13~0.98 | 0.43 | 0.64 |

In 2017, the experimental data is measured 50 groups of which 25 groups are used to build the model of rice roots and others are used to test the model. In order to test the accuracy of the morphological size of rice roots using the root model, the average values of running 25 times simulations are computed and compared with the experimental data of rice roots collected in 2016. The measured values and simulation values of the total root length, the total root area, and the Total root volume of single plant are compared respectively. The comparison results are shown in Figure 3. The experimental results demonstrate that the values of the total root length, the total root area, and the total root volume of single plant are almost corresponded with the measured values achieved by the model. Therefore, the above model of rice roots can better predict the dynamic changes of the roots with the root length, the root area and the root volume during rice growth under the different conditions. The results of test show that the root model achieves the good reliability.

**Fig.3.**
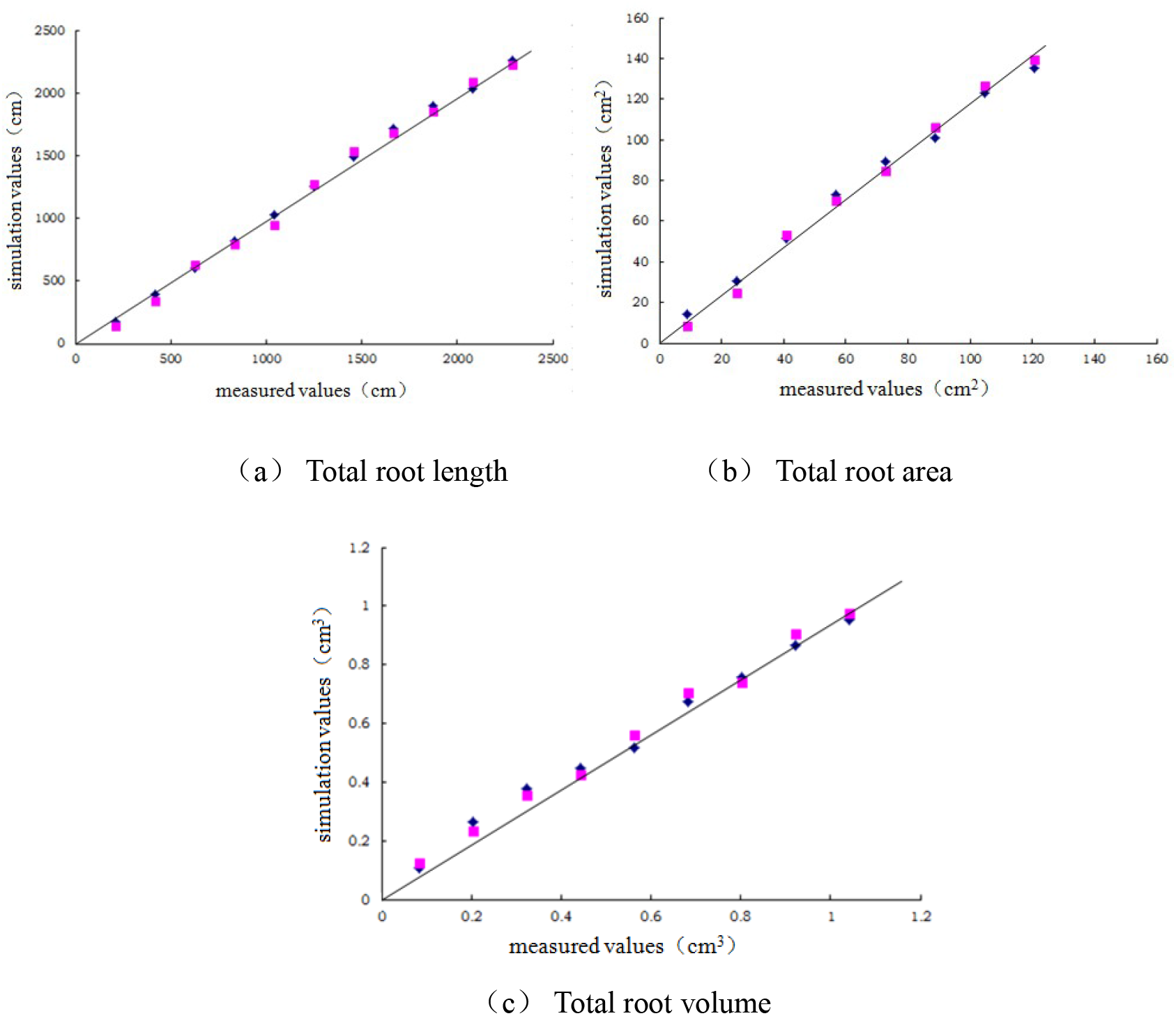
The comparison of observed values and simulated values

### 4.2 Comparison of different simulation solutions

According to the measured data and the growth trace of rice root morphological index and taking single rice roots as an example, in 3D space, the simulation of single rice roots is done using three different simulation solutions respectively, which the results is shown in Figure 4.

**Fig.4.**
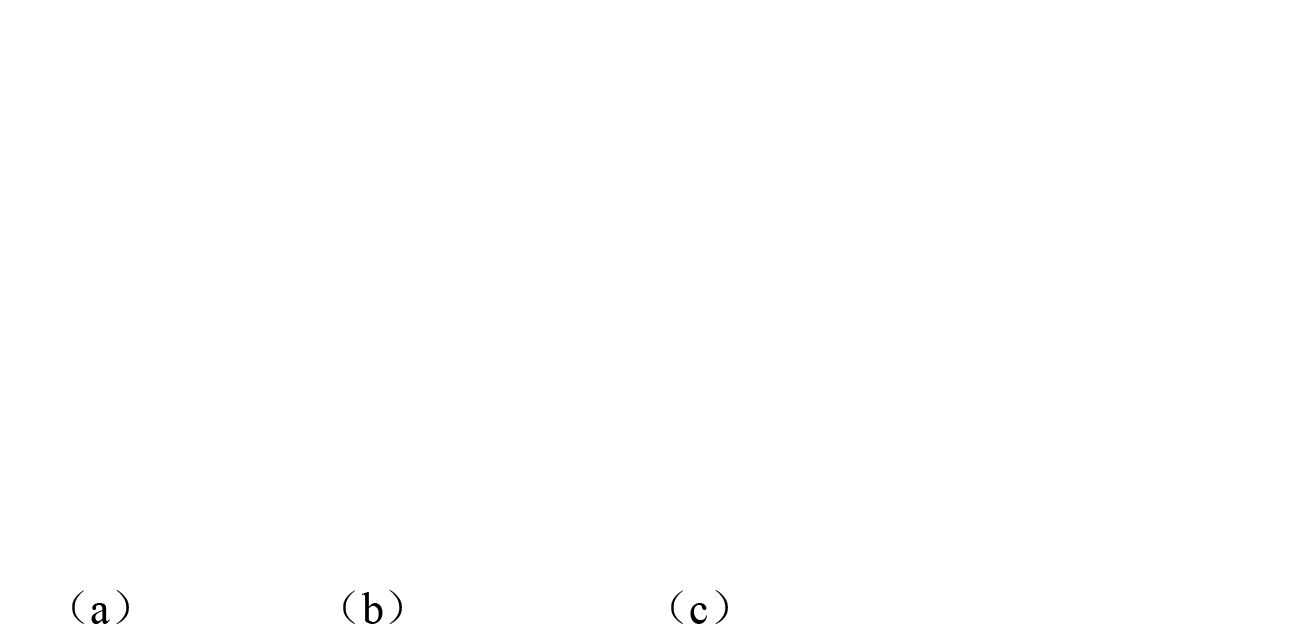
Schematic diagram of Single Rice Root Morphology Simulation

In Figure 4(a), the rice root simulation results are shown by L-system iterating twice to generate. From it we can see that it is not ideal for the curve degree because the main root and branch roots of levels of roots are straighter. Moreover, the surface of roots exists the wrinkle, which can only be used to barely express the topological structure of rice roots. The Figure 4(b) is the rice root simulation chart which the improved L-system generated. The system is combined with B-spline curve to optimize deeply the 3D display effect of root images. The results show that the generation of roots is more closed to smooth and the combination of the erect and curve. It can express the curve of rice root geometric morphology.It is more natural for the root morphology. However, it is not strong for the three-dimension effect of the generating map of the whole root. Therefore, around the rice root growth trace, the terraces are adopted to draw the root segments and all root segments are connected together to form a whole root, which generates the rice root simulation chart in Figure 4(c). As can be seen from Figure 4(c), the chart presents the smoothness and convexity and it can better reflect the natural smoothness and overall fidelity of rice roots.

### 4.3 The 3D visualization simulation of rice roots

According to the 3D display model of single root rice roots and the rice roots model based on morphological parameters and the topological structure base on the rice roots, the 3D visualization simulation of rice roots is constructed by using the Visual C++ operation platform and combining OpenGL standard graphic library. In the system, the angle between roots, the curve degree of branch roots and the extension situation of root growth should be considered. The system adopts random functions to describe the possible random changes of various quantities, and controls the growth of rice roots by controlling the diameter, length and iteration number of root nodes. The iteration number is mainly used to control the topological structure of roots. So the more the number of iterations, the denser the rice roots become. The growth of diameter and length causes the roots to grow up continuously and produces the effect of root growth. In this paper, a hybrid rice variety Ganxin 203 is used as a test object. According to the description of the morphology and growth characteristics of rice root growth at different development stages, the L-system production formula of root characteristics was expressed by combined with Cubic growth function and B-spline curve. The three-dimensional image of rice roots was real-time processed by color rendering, illumination processing and texture mapping. The dynamic growth process of rice roots is illustrated in Figure 5.

**Fig.5.**
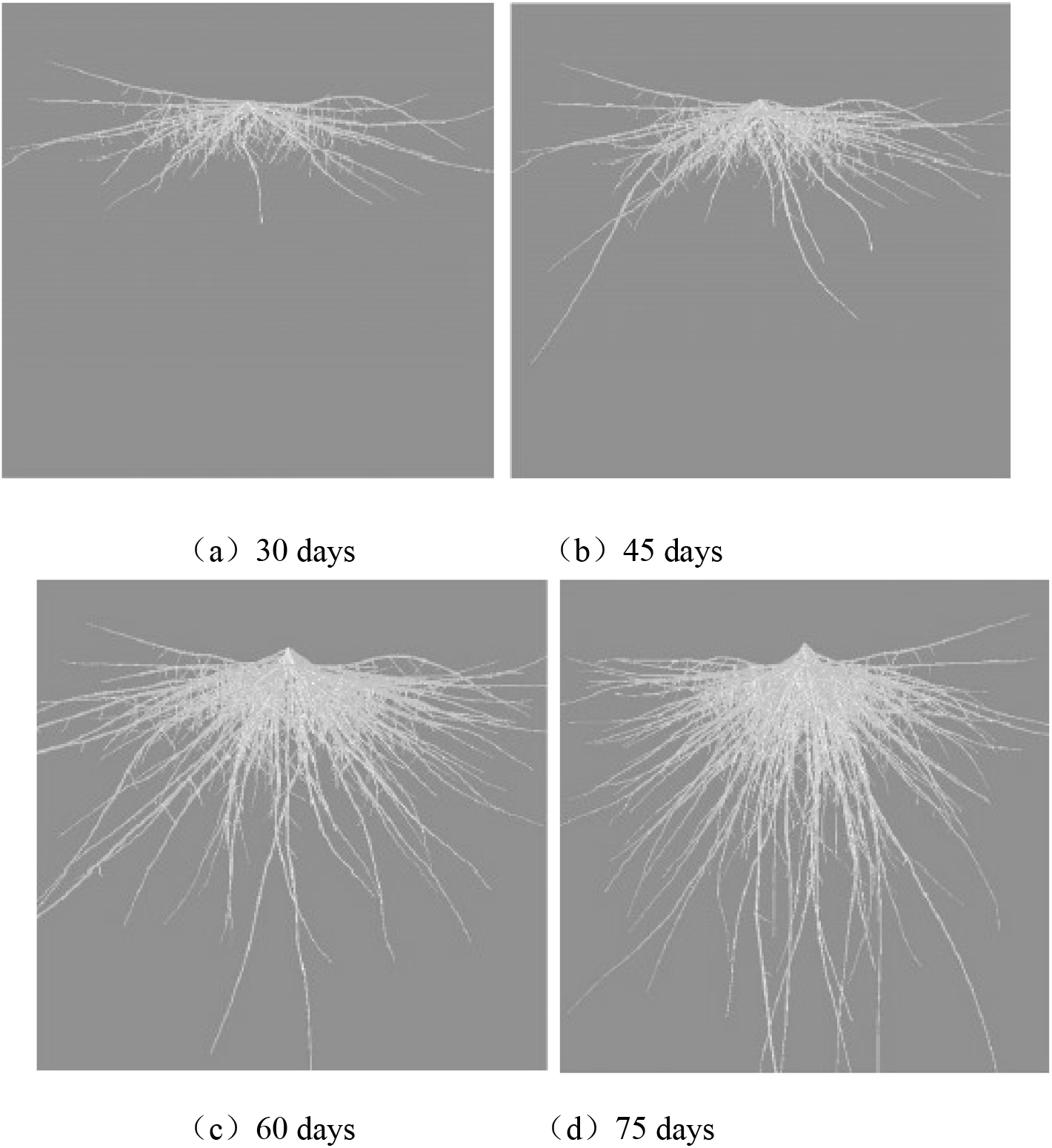
Diagram of the dynamic model of rice roots

## 5 Conclusions

The modeling and visualization simulation of rice roots is an evidently important part for constructing virtual rice. This paper adopts the “root box” experiment to observe and analyze the spatial morphological distributional characteristics of rice roots and studies the morphological structure, the growth and distributional characteristics during the growth and development of rice roots. Meanwhile, in this paper, the L-system of rice roots is designed, and the rice root model based on morphological parameters is constructed by optimizing the Cubic growth function and combining with B-spline curve.The visualization simulation of rice roots is also implemented by using the computer graphic technology. The results show that the model simulates the geometric morphological parameter to show their growth and development law preferably. The data achieved by the model is better corresponded with the measured data.The correlation coefficient is 0.997 to run up to an extremely significant level with a smaller coefficient variation. From the 3D images of the system we can see they are very similar with ones of real rice roots in the morphological structure, which indicates that it is very effect to use the root model to simulate the growth and development of rice roots. But in this experiment many environmental factors such as air, water, nitrogen and light are ignored. Moreover, the factors such as the errors collecting the experimental data of rice roots and limitations causes the experimental data to deviate greatly from the real data in natural environment. Therefore, in the future work, the greenhouses based on the field-bus control system are used to cultivate rice. The differential L-system is used to simulate the discrete continuous situation of rice root growth. Meanwhile further combined with physiological and ecological factors the visualization simulation is implemented to obtain more realistic simulation effect, which makes the simulation images closer to the natural ones.

## Acknowledgment

This work is supported by the National Natural Science Foundation of China (No. 61862032), the Science and Technology Plan Projects of Jiangxi Provincial Education Department (No. GJJ160409).

